# Rem2 interacts with CaMKII at synapses and restricts long-term potentiation in hippocampus

**DOI:** 10.1101/2024.03.11.584540

**Authors:** Rabia Anjum, Vernon R J Clarke, Yutaro Nagasawa, Hideji Murakoshi, Suzanne Paradis

## Abstract

Synaptic plasticity, the process whereby neuronal connections are either strengthened or weakened in response to stereotyped forms of stimulation, is widely believed to represent the molecular mechanism that underlies learning and memory. The holoenzyme CaMKII plays a well-established and critical role in the induction of a variety of forms of synaptic plasticity such as long-term potentiation (LTP), long-term depression (LTD) and depotentiation. Previously, we identified the GTPase Rem2 as a potent, endogenous inhibitor of CaMKII. Here, we report that knock out of *Rem2* enhances LTP at the Schaffer collateral to CA1 synapse in hippocampus, consistent with an inhibitory action of Rem2 on CaMKII in vivo. Further, re-expression of WT Rem2 rescues the enhanced LTP observed in slices obtained from *Rem2* conditional knock out (cKO) mice, while expression of a mutant Rem2 construct that is unable to inhibit CaMKII in vitro fails to rescue increased LTP. In addition, we demonstrate that CaMKII and Rem2 interact in dendritic spines using a 2pFLIM-FRET approach. Taken together, our data lead us to propose that Rem2 serves as a brake on runaway synaptic potentiation via inhibition of CaMKII activity. Further, the enhanced LTP phenotype we observe in *Rem2* cKO slices reveals a previously unknown role for Rem2 in the negative regulation of CaMKII function.

## Introduction

CaMKII is an abundant, multifunctional serine-threonine kinase whose regulation and activity is uniquely sensitive to synaptic activity [1]. Decades of biochemical, genetic, and electrophysiological experiments established a critical role for CaMKII in the synaptic strengthening (i.e. LTP) that underlies learning and memory in the hippocampus [2,3]. CaMKII monomers contain an N-terminal catalytic domain, central regulatory domain, and a C-terminal association domain, which mediate assembly into the dodecamer, holoenzyme form [4]. In the canonical pathway for LTP induction at CA3-CA1 synapses of the hippocampus, high frequency stimulation triggers the activation of NMDARs [5]. Subsequent Ca^2+^ influx through the NMDAR ionophore activates calmodulin (CaM) which binds to the holoenzyme CaMKII causing a conformational change, displacing the regulatory domain and exposing the substrate binding site(s) and, notably, a threonine residue (T286) on the regulatory domain [6]. Trans-autophosphorylation of the T286 site by an activated CaMKII catalytic domain results in an autonomously active form of CaMKII which allows its activity to persist even in the absence of Ca^2+^ [7]. LTP is expressed as an increase in AMPAR function: increased number of receptors [8] and increased single channel conductance [9].

Many studies point to a role for the catalytic activity of CaMKII in LTP expression via phosphorylation of downstream targets (e.g. the GluA1 subunit of the AMPA receptor) subsequent to T286 phosphorylation [10–13]. Manipulations that block the catalytic activity of CaMKII such as application of peptide inhibitors [2,14] or deletion of the CaMKIIα gene impair LTP [15]. However, it is also appreciated that CaMKII binding to the GluN2B subunit of the NMDAR [16,17] plays a critical role in LTP induction [15,18,19]. In fact two recent studies demonstrate that the enzymatic function of CaMKII towards non-CaMKII substrates is dispensable for LTP induction while CaMKII binding to GluN2B, a “structural” function of CaMKII, is required [20,21]. In addition, new x-ray crystallographic structures reveal that CaMKII substrates bind along one continuous binding site [22] opposed to two distinct sites (i.e. S- and T-sites) on the catalytic domain [23]. Because inhibitors of CaMKII catalytic activity which are known to impair LTP also block GluN2B binding [14] the critical role of CaMKII-GluN2B binding in LTP was obscure until now.

The GTPase Rem2 is a potent (K_i_ ∼6nM) inhibitor of CaMKII catalytic activity *in vitro* (ROYER). Rem2 inhibits the autonomously active CaMKII holoenzyme, does not interfere with CaMKII autophosphorylation at T286, and does not interfere with CaM binding [24]. Based on these data we favor a model by which the N-terminus of Rem2, which contains a CaMKII pseudosubstrate sequence, inhibits CaMKII in vitro by binding to the substrate binding site in the CaMKII catalytic domain. In so doing, Rem2 may also block access of the GluN2B binding site on CaMKII.

Rem2 is a member of the RGK family of non-canonical Ras-like GTPases (which also includes Rad, Gem/Kir, and Rem) and is primarily expressed in the brain [25]. A number of features differentiate RGK family members from the Ras superfamily. For example, the crystal structures of several RGK proteins, including Rem2, reveal differences between this family and classical GTPases in the structure of their nucleotide binding domains [26,27]. For the most part, neither GTPase activating proteins (GAPs) nor guanine nucleotide exchange factors (GEFs) have been identified for this family [28] (the only exception being the identification of NmeI as a Rad GAP [29]). For these and other reasons, Rem2 and the RGK family members in general may not behave as classical Ras-like GTPases regulated by their nucleotide binding state [30].

Overexpression of RGK family members in variety of cell types results in cytoskeleton rearrangements and decreased Ca^2+^ influx through voltage-gated calcium channels [31–35] suggesting that RGK proteins are multifunctional signal transducers. However, the signaling mechanism(s) by which Rem2 and other RGK proteins mediate these processes are not well understood. Previous studies from our lab using *Rem2* knockout approaches demonstrated that Rem2 is a critical regulator of dendritic branching, synapse formation and cortical plasticity in the vertebrate nervous system [35–41]. In addition, Rem2 and CaMKII co-associate in neurons [24,42] and interact in a signal transduction pathway that regulates dendritic arborization [24,37,38].

Here we test the hypothesis that Rem2 plays a previously unsuspected role in CaMKII-dependent LTP. We posited that Rem2 could influence synaptic plasticity through an interaction with CaMKII in the postsynaptic dendritic spine, the site of CaMKII activation and LTP induction and expression. To test this hypothesis, we studied the expression of Rem2 in hippocampus. We induced LTP at the Schaffer collateral to CA1 (SC-CA1) synapse in acute hippocampal slice isolated from *Rem2* WT or *Rem2* cKO mice using high frequency stimulation and found the loss of Rem2 was associated with increased levels of synaptic potentiation. Further, the effect of Rem2 on LTP was dependent on CaMKII. Finally, we demonstrated the ability of Rem2 and CaMKII to interact in dendritic spines in response to glutamate stimulation using 2pFLIM-FRET and glutamate uncaging in organotypic hippocampal slice preparations.

## Methods

### Animals

For LTP experiments, *Rem2^flx/flx^* mice [40] were bred and housed in the animal facility at Brandeis University. For FRET-FLIM experiments C57BL/6N mice were purchased from Japan SLC, Inc. and bred and housed in the animal facility in the National Institute of Natural Sciences. In both cases, animals were maintained with a 12-hour light-dark cycle. Food and water were available *ad libitum*. All animal procedures were approved by the Brandeis University Institutional Animal Care and Usage Committee or National Institute of Natural Sciences Animal Care and Use Committee, and all experiments were performed in accordance with relevant guidelines and regulations.

### Reagents

4-methoxy-7-nitroindolinyl-caged-L-glutamate (MNI-caged glutamate) was purchased from Tocris Bioscience (1490). Myristoylated CaMKIINtide, i.e. myr-CN27, was purchased from EMD Millipore (208921).

### Plasmids

Plasmids containing CaMKIIα, CaMKII promoter with 0.4 kbp (CaMKII0.4), WPRE genes were gifts from Y. Hayashi, M. Ehlers, and K. Kobayashi, respectively. Plasmids containing Clover, WPRE3, Cre, and hSyn-DIO-EGFP genes were gifts from M. Lin, Bong-Kiun Kaang, Connie Cepko, and Bryan Roth (Addgene plasmid #40255, #61463, #13775, #50457), respectively. pAAV-CaMKII0.4-DIO-Clover-CaMKIIα-WPRE3, pAAV-CaMKII0.4-DIO-mCherry-Rem2-WPRE3, and hSyn-Cre-WPRE plasmids were constructed by inserting the respective components into the pAAV-MCS.

### Virus injections

AAV-GFP (pAAV9-hSyn-EGFP; 50465-AAV9, 6.33 × 10^12^ vg/mL), AAV-GFP-Cre (pENN.AAV9.hSyn.HI.eGFP-Cre.WPRE.SV40, 105540-AAV9, 2.2 × 10^12^ vg/mL), AAV-tdtomato (pAAV9-CAG-tdTomato; 59462-AAV9, 8.33 × 10^12^ vg/mL), AAV-tdtomato-flex (pAAV9-FLEX-tdTomato; 28306-AAV9, 1.03 × 10^13^ vg/mL), and AAV-mCherry (pAAV9-hSyn-mCherry; 114472-AAV9, 9.0 × 10^12^ vg/mL) viruses were purchased from Addgene. AAV-Rem2-wt (AAV9-hSyn1-Myc-Rem2-WPRE, 3.3 × 10^12^ GC/ml) and AAV-Rem2^RR/GG^ (AAV9-hSyn1-Myc-Rem2^RR/GG^-WPRE, 3.3 × 10^12^ GC/ml) viruses were produced and purified by Vector Biolabs. Mice between P26-P30 were anesthetized and mounted in a stereotaxic frame and the skull exposed. To target the dorsal CA1 of the hippocampus, the coordinates used were as follows: A/P −1.95 mm, M/L ± 1.40 mm, and D/V 1.15 mm. A small 1–2 mm in diameter hole was drilled in the skull in both hemispheres, and a glass micropipette delivered 400 nl of virus in each hemisphere. Animals were allowed to recover for 2 weeks before experiments were performed.

### Immunohistochemistry in slices

*Rem2^flx/flx^* mice were injected with AAV-GFP-Cre, AAV-Rem2-wt and AAV-Rem2^RR/GG^ at P27-P28 as described above. At 14 dpi, transcardial perfusion was performed with 4% paraformaldehyde/4% sucrose in 1X PBS solution. The brains were removed and fixed in 4% paraformaldehyde/4% sucrose for 24 h and then transferred to 30% sucrose for 24 h for storage. The brains were sectioned at 50 µm using a microtome (Leica) and slices were suspended in PBS at 4°C until being processed for immunohistochemistry.

Brain sections containing hippocampus were blocked in PBS with 20% BSA and 0.1% Triton for two hours at room temperature prior to incubation with the primary antibody mouse anti-Rem2 (Santa Cruz, sc-514999) at a concentration of 1:100 overnight at 4°C. The sections were then incubated in the secondary antibody (goat anti-mouse Alexa Fluor™ 546, Thermofisher A-21052 or goat anti-mouse Alexa Fluor™ 633, Thermofisher A-21052) at 1:500 for two hours at room temperature and mounted with Vectashield Hardset antifade mounting medium with DAPI (Vector Laboratories, H-1500).

### Image acquisition and analysis

Images were acquired on a Zeiss LSM880 confocal microscope using the Plan-Apochromat 63x/1.40 Oil DIC M27 objective. We acquired z-stacks (1.0 µm step size) of DAPI-positive and (where applicable) Cre-positive cell somas located within the pyramidal cell layer region of the CA1 subregion of the dorsal hippocampus. For each section, images including the pyramidal cell layer and *striatum oriens* were stitched together (∼600 µm M/L and ∼200 µm D/V) using the tile scan function; 2-3 sections were imaged per animal. Microscope settings for laser power, detector gain, and amplifier offset were initially optimized across multiple no injection control coverslips and were kept constant within each experimental round.

In order to minimize background noise, maximum intensity projection images were pre-processed in the following manner using ImageJ. In the channel containing anti-Rem2 staining, a small region of interest (∼75×75 µm) was drawn in an area where neither cell bodies nor anti-Rem2 staining was visible, and the background in the anti-Rem2 channel was measured using the Analyze > Measure menu option in ImageJ. Then, the background was subtracted from the anti-Rem2 channel using the Process > Math > Subtract menu option, using the background value measured in the previous step.

Cell detection and counting and subsequent quantification of anti-Rem2 staining was carried out using a semi-automatic pipeline in QuPath with maximum intensity projection images. Using the DAPI channel, a region of interest in the CA1 area of the hippocampus was drawn around the pyramidal cell layer in each stitched image. Within this region of interest, neuronal somas and nuclei were detected based on the DAPI-stained nucleus using the Analyze > Cell Detection menu option in QuPath, with software recommended parameters. Rem2+ staining in the cell soma was detected by creating a classification filter, using the Classify > Object classification > Create single measurement classifier menu option in QuPath. A fixed pixel threshold value was empirically determined for mean fluorescence in the cytoplasm in the α-Rem2 channel. This threshold value was kept constant and applied to all images within each experimental round.

Quantification of anti-Rem2 staining in the dendritic arbors of CA1 pyramidal neurons was carried out using ImageJ with background subtracted maximum intensity projection images. A rectangular region of interest (∼150×100 µm) was drawn in the *striatum radiatum* within the CA1 of the hippocampus in each stitched image. The average pixel intensity in the channel for anti-Rem2 staining was measured using the Analyze > Measure > Mean Gray Value menu option in ImageJ.

### Field electrophysiology recordings

Brains were isolated from *Rem2^flx/flx^* mice of both sexes 12-16 days post-injection (d.p.i). The brain was rapidly dissected and placed in ice-chilled, oxygenated modified artificial cerebrospinal fluid (aCSF) comprising of (in mM): NaCl 124, KCl 3.7, NaHCO_3_ 24.6, CaCl_2_ 1, MgSO_4_ 4, D-glucose 10, KH_2_PO_4_ 1.2 saturated with 95% O_2_ / 5% CO_2_. Parasagittal slices (400 µm thick) containing the hippocampal region were prepared in ice-chilled, oxygenated aCSF using a Leica VT1200 vibrating microtome (Leica Biosystems Inc.).

Following equilibration for at least an hour, slices were transferred to a brain slice interface chamber (model BSC2, Scientific Systems Design, Inc.); slices rested on filter paper at the interface of the perfusing solution (0.4-0.6 ml min^−1^) comprising of standard aCSF (in mM): NaCl 124, KCl 3.7, NaHCO_3_ 24.6, CaCl_2_ 2.5, MgSO_4_ 1, D-glucose 10, KH_2_PO_4_ 1.2 saturated with 95% O_2_ / 5% CO_2_. Field recordings were obtained using a glass microelectrode (resistance ∼ 2–4 MΩ) containing 3 M NaCl or standard aCSF, placed at the radiatum border. Responses were evoked by stimulating two electrodes placed within the Schaffer-collateral-commissural fibers. Baseline stimulation was every 30 s for each input with a 15 s interval between alternating inputs. Independence of inputs was assessed using a paired-pulse protocol. Stimulation intensity was adjusted such that the fEPSP slope was approximately 20-40% of that required to generate a visible population spike. LTP was induced by tetanic stimulation (100 Hz; 1 s) delivered 3 times at 5 minute intervals. The myristoylated CaMKIINtide, myr-CN27 (EMD Millipore, 208921) was bath applied where indicated. Recordings were filtered at 3–10 kHz using an Axoclamp 2A Amplifier (Molecular Devices), and collected for online analysis at a sampling rate of 20 kHz using WinLTP software [43].

### Organotypic hippocampal slice culture and virus infection

Hippocampal slices were prepared from P6-P9 C57BL/6N mice as described previously [44]. Briefly, the animal was deeply anesthetized with isoflurane, after which the animal was quickly decapitated, and the brain removed. The hippocampi were isolated and cut into 350 µm sections in an ice-cold dissection medium (250 mM N-2-hydroxyethylpiperazine-N′-2-ethanesulfonic acid, 2 mM NaHCO_3_, 4 mM KCl, 5 mM MgCI_2_, 1 mM CaCl_2_, 10 mM D-glucose, and 248 mM sucrose). The slices were cultured on the membrane inserts (PICM0RG50; Merck, Ireland), placed on the culture medium (50% minimal essential medium, 21% Hank’s balanced salt solution, 15 mM NaHCO_3_, 6.25 mM N-2-hydroxyethylpiperazine-N′-2-ethanesulfonic acid, 10 mM D-glucose, 1 mM L-glutamine, 0.88 mM ascorbic acid, 1 mg/mL insulin, and 25% horse serum), and incubated at 35°C in 5% CO_2_.

The cultured slices at DIV3 were transfected with AAV using a glass pipette (Narishige). The preparation of AAV (titers typically ∼5 × 10^9^ genome copies/µl) has been described previously in detail [45,46]. For sparse labeling, AAVs encoding Clover-CaMKIIα and mCherry-Rem2 were coinfected with a low amount of AAV encoding Cre with the ratio of 100:300:4. On DIV13 or 14, two-photon FLIM-FRET experiment was carried out.

### Two-photon fluorescence lifetime imaging and MNI-glutamate uncaging

Details of two-photon FLIM-FRET imaging were described previously [47,48]. Briefly, Clover-CaMKIIα was excited at 1000 nm with a Ti: sapphire laser (Mai Tai; Spectra-Physics). The scanning mirrors were controlled with the ScanImage software [49]. The Clover fluorescence was collected by an objective lens (60×, 1.0 NA; Olympus) and a photomultiplier tube (H7422-40p; Hamamatsu) placed after a dichroic mirror (565DCLP; Chroma) and emission filter (FF01-510/84; Semrock).

Measurement of fluorescence lifetime was conducted using a time-correlated single-photon counting board (SPC-150, Becker & Hickl GmbH) controlled with custom software [47]. The fluorescence lifetime image was constructed by translating the mean fluorescence lifetime in each pixel into a color-coded image [48]. Analysis of the lifetime change and binding-fraction change was conducted as described elsewhere [48].

Two-photon MNI-glutamate uncaging was carried out in the imaging buffer solution (136 mM NaCl, 5 mM KCl, 0.8 mM KH_2_PO_4_, 20 mM NaHCO_3_, 1.3 mM L-glutamine, 0.2 mM ascorbic acid, MEM amino acids solution [Thermo Fisher], MEM vitamin solution [Thermo Fisher], and 1.5 mg/mL phenol red) containing 4 mM CaCl2, 0 mM MgCl2, 1 mM tetrodotoxin, and 1 mM MNI-caged glutamate aerated with 95% O_2_/5% CO2 at 24–26 ℃. Glutamate uncaging was carried out at 720 nm (30 trains, 0.5 Hz, 6 ms duration/pulse, 6 mW) at the tip of the spines.

### Statistical analysis

Statistical analyses were performed using the R programming language [24]. For all experiments we assessed the basic properties of samples (Q-Q plots, residual plots, distributions etc.) prior to running statistical tests with distribution assumptions. LTP experiments were analyzed using nonparametric bootstrapped multiple comparisons tests using custom written code in R. The *p* values were obtained were controlled using the Sidak-Holm correction. Box plots are used to graphically represent summary data: the center line is the median, the edges of the box mark the interquartiles of the distribution and the lines extend to the limiting values of the sample distribution when outliers are excluded (defined as values further away from the median than 1.5 × interquartile range)

## Results

Rem2 is expressed almost exclusively in the nervous system [25] and previous studies from our lab demonstrate that Rem2 is expressed in cortex and hippocampus [37,40]. In addition, we showed that Rem2 mRNA expression is up-regulated in response to neuronal depolarization in cultured cortical neurons [39] as well as in response to sensory experience in visual cortex in the intact mouse brain [40]. In hippocampus, Rem2 is expressed predominantly in the pyramidal cell layer of the CA1-CA3 subregions in young adult animals [50,51].

Given the expression of Rem2 in pyramidal neurons in CA1 area of hippocampus, we sought to determine if Rem2 plays a role in CaMKII-dependent LTP at the CA3-CA1 Schaffer collateral synapse. To restrict Rem2 deletion to the CA1 region of the hippocampus, we injected Cre-expressing virus into that region in a *Rem2^flx/flx^* mouse [40] by stereotaxis. Specifically, we injected an AAV9-hSyn-CreGFP virus by itself (*Rem2* cKO) or we coinjected with a virus expressing either wild-type Rem2 (*Rem2* cKO + wt-Rem2) or a mutant Rem2 which renders it unable to inhibit CaMKII kinase activity (*Rem2* cKO + Rem2^RR/GG^) [24].

In order to validate *Rem2* deletion and expression of rescue constructs, we quantified Cre-GFP-positive and Rem2-positive cell bodies across conditions using an antibody that recognizes the Rem2 protein (Figure 2A). We found that Cre-GFP was detected in the majority of cell bodies in all three experimental conditions (*Rem2* cKO, 96.5 ± 0.3%, 221 ± 17); *Rem2* cKO + wt-Rem2, 92.5 ± 1.1%, 175 ± 15; *Rem2* cKO + Rem2^RR/GG^; 89.0 ± 0.7%, 185 ± 17), indicating that virus was delivered to the targeted area (Figure 2B, C). We also detected Rem2 immunostaining in the majority of cell bodies in the conditions in which the Rem2 protein is re-introduced by viral mediated gene transduction (*Rem2* cKO + wt-Rem2, 73.5 ± 0.5%, 136 ± 10; *Rem2* cKO + Rem2^RR/GG^; 97.0 ± 0.0%, 201 ± 20) with minimal staining in sections isolated from *Rem2* cKO animals as expected (*Rem2* cKO, 12.5 ± 1.5%, 31 ± 14). In addition, GFP signal and anti-Rem2 staining was detected in the majority of neurons in CA1 in the animals that were injected with virus expressing either wt-Rem2 or Rem2^RR/GG^ and Cre-GFP (Figure 2 A-C) (*Rem2* cKO + wt-Rem2, 70.0 ± 1.0%, 130 ± 11; *Rem2* cKO + Rem2^RR/GG^; 87.5 ± 0.5%, 182 ± 17), demonstrating that we achieved co-infection of viruses in a high percentage of neurons in the targeted area.

**Figure 1.**
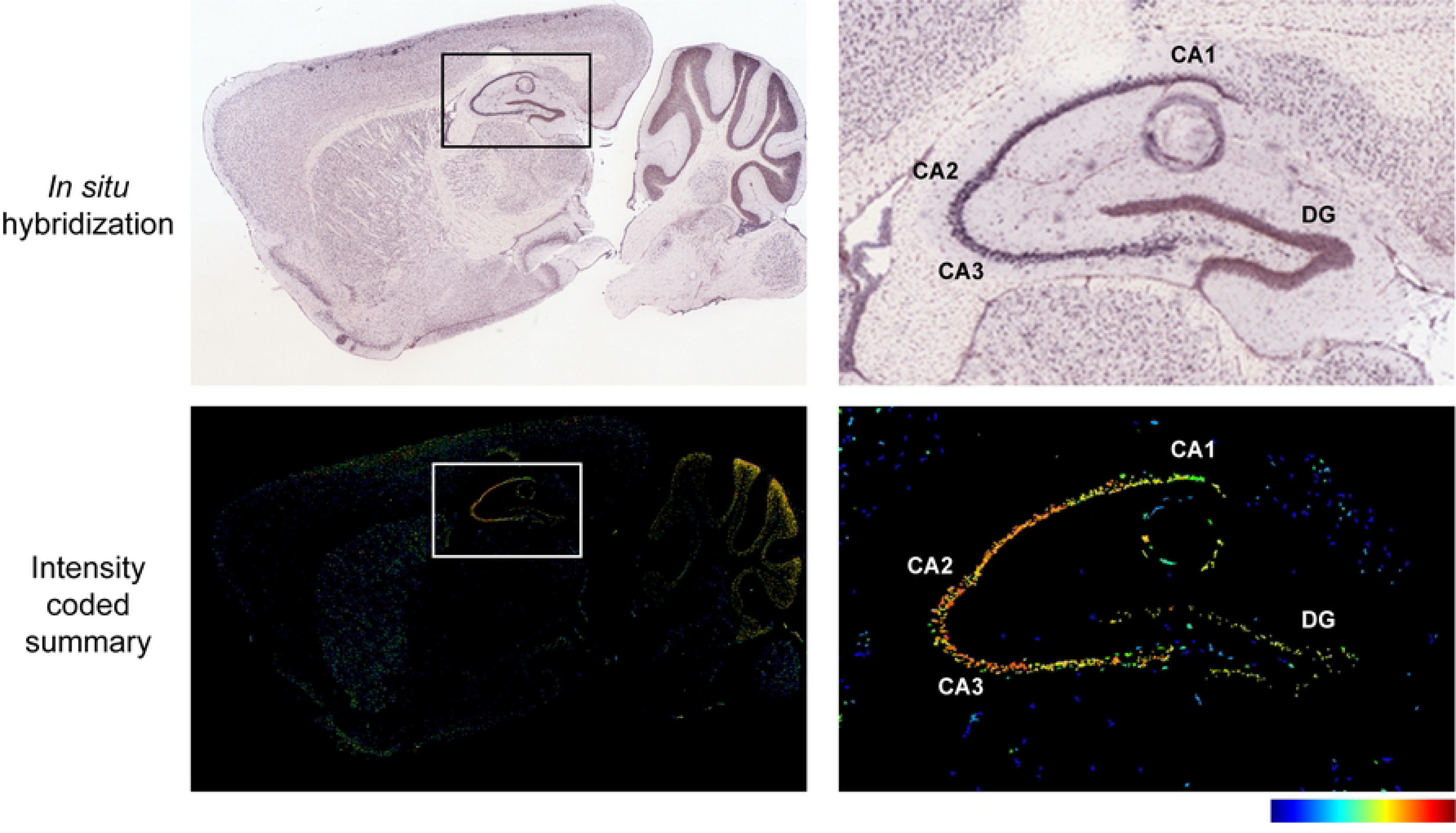
Rem2 mRNA is expressed in the hippocampus. (A-B) In situ hybridization (ISH) images of *Rem2* mRNA in sagittal brain sections from P56 mice. (C-D) Intensity-coded summary images show low (blue) to high (red) Rem2 expression. Box (A and C) indicates the hippocampus, which is expanded on the right (B and D). Image credit: Allen Mouse Brain Atlas, https://mouse.brain-map.org/gene/show/80094.

**Figure 2.**
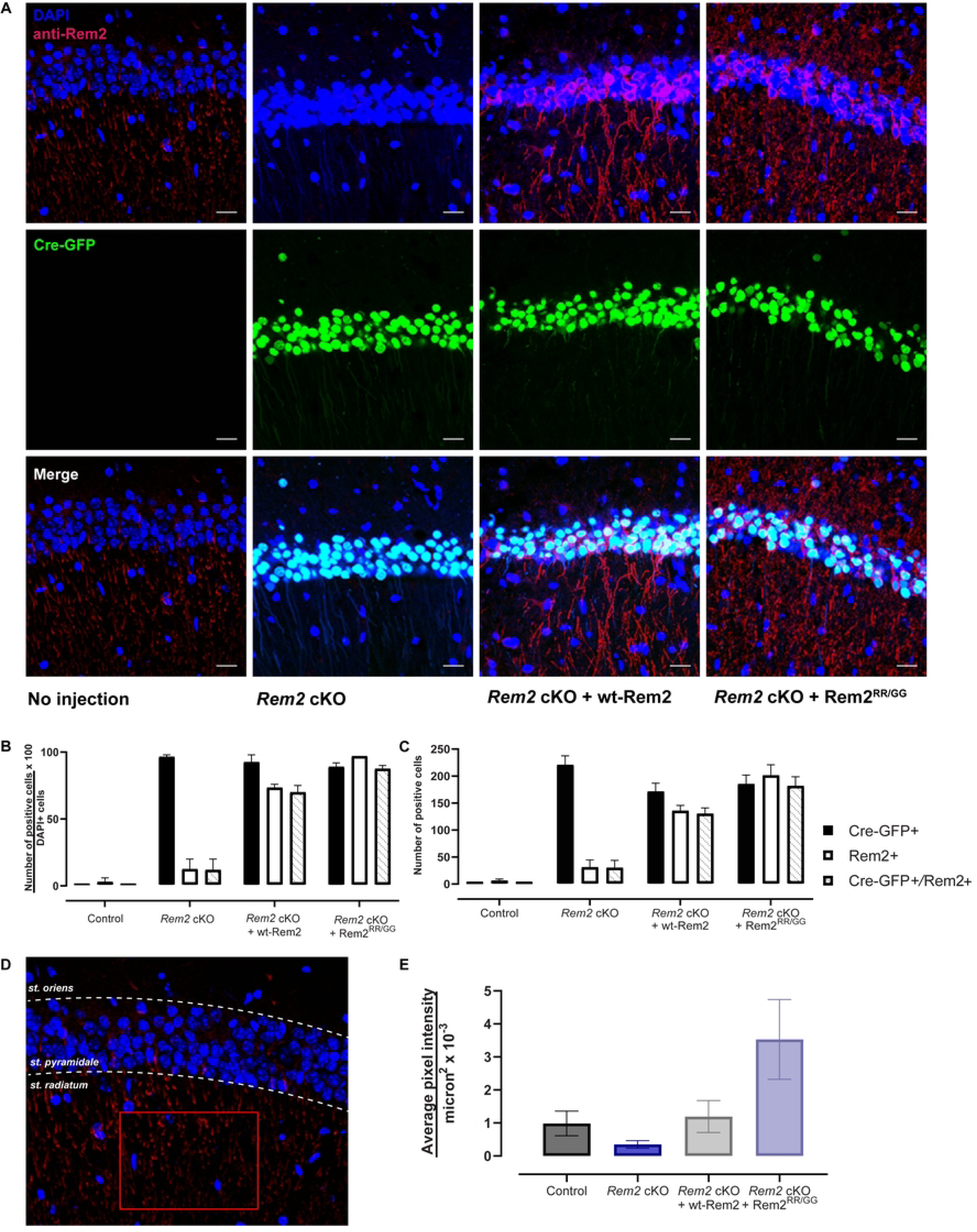
Viral-mediated conditional knockout of *Rem2* and rescue with either wt-Rem2 or Rem2^RR/GG^ in hippocampus from P28 mice. A) Representative images from coronal sections of the dorsal CA1 from uninjected and virus-injected mice. Top, DAPI and anti-Rem2; middle, Cre-GFP; bottom, merged image. DAPI = blue, Cre-GFP = green, anti-Rem2 = red. B) Quantification of the percentage of neurons expressing indicated markers divided by DAPI-positive neurons for each experimental condition. C) Total number of neurons expressing indicated markers for each condition. D) Schematic for quantifying anti-Rem2 staining in the dendritic arbor of pyramidal neurons, from an uninjected control animal. A region of interest was drawn (red rectangle) in the striatum radiatum within the CA1 region of the hippocampus, the average pixel intensity in the channel for anti-Rem2 staining was measured. E) Average pixel intensity per square microns for each condition. Error bars = SEM. 350-1000 cells per condition from 2-3 slices per biological replicate in 2 biological replicates. Scale bar = 20 µm. st. = striatum.

Somewhat unexpectedly we were unable to detect appreciable anti-Rem2 staining in the soma of the uninjected (i.e. wild-type) control animals (Figure 2B and 2C). However, we observed anti-Rem2 staining in the dendritic arbor of pyramidal neurons in the uninjected controls, suggesting that perhaps Rem2 protein is more abundant in cell processes in these P42 animals. To quantify this expression, we drew a region of interest in the striatum radiatum of the CA1 region (Figure 2D) and measured the average pixel intensity per square micron in the anti-Rem2 channel. Our analysis shows that anti-Rem2 staining in striatum radiatum is observable in both uninjected control and animals injected with Cre-GFP and a virus expressing wt-Rem2 at a similar level (Control 0.985 ± 0.373 per µm^2^ × 10^−3^; *Rem2* cKO + wt-Rem2 1.193 ± 0.485 per µm^2^ × 10^−3^). Animals injected with Cre-GFP and a virus expressing Rem2^RR/GG^ showed a higher level of anti-Rem2 staining (*Rem2* cKO + Rem2^RR/GG^ 3.529 ± 1.208 per µm^2^ × 10^−3^), while animals injected with Cre-GFP alone showed minimal anti-Rem2 staining (*Rem2* cKO 0.348 ± 0.284 per µm^2^ × 10^−3^) confirming that the antibody detects Rem2 expression in all of our samples including uninjected controls.

Next, we sought to determine if *Rem2* deletion affects long-term potentiation. We injected 4-week-old *Rem2^flx/flx^* mice with an AAV9-hSyn-GFP (control) or AAV9-hSyn-CreGFP virus (*Rem2* cKO), or AAV9-hSyn-CreGFP together with a virus expressing wt-Rem2 (*Rem2* cKO + wt-Rem2), or AAV9-hSyn-CreGFP together with a virus expressing Rem2^RR/GG^ (*Rem2* cKO + Rem2^RR/GG^) into the CA1 region of the hippocampus. We subsequently performed field recordings from the CA1 area of acute hippocampal slices isolated from animals 14 days post-injection (d.p.i.). We induced LTP at SC-CA1 synapse using high frequency stimulation (comprising 3 tetanic stimulations of 100Hz for 1s delivered at intervals of 5 minutes) at *t =*30 in control slices (Figure 3A, E; *n =* 10). Our results show that *Rem2* deletion enhances potentiation compared to control slices (Figure 3B, E; *n =* 10), which lasted for at least 130 minutes post-tetanus. Additionally, while re-expression of wt-Rem2 rescues the enhanced LTP phenotype (Figure 3C, E; *n =* 10), expression of Rem2^RR/GG^ fails to rescue the enhanced LTP that is observed upon *Rem2* knockout (Figure 3D, E; *n =* 10). From these data we conclude that Rem2 expressed in pyramidal neurons in CA1 normally acts as a brake on LTP at this synapse and further, that Rem2 inhibition of CaMKII is crucial to this process.

**Figure 3.**
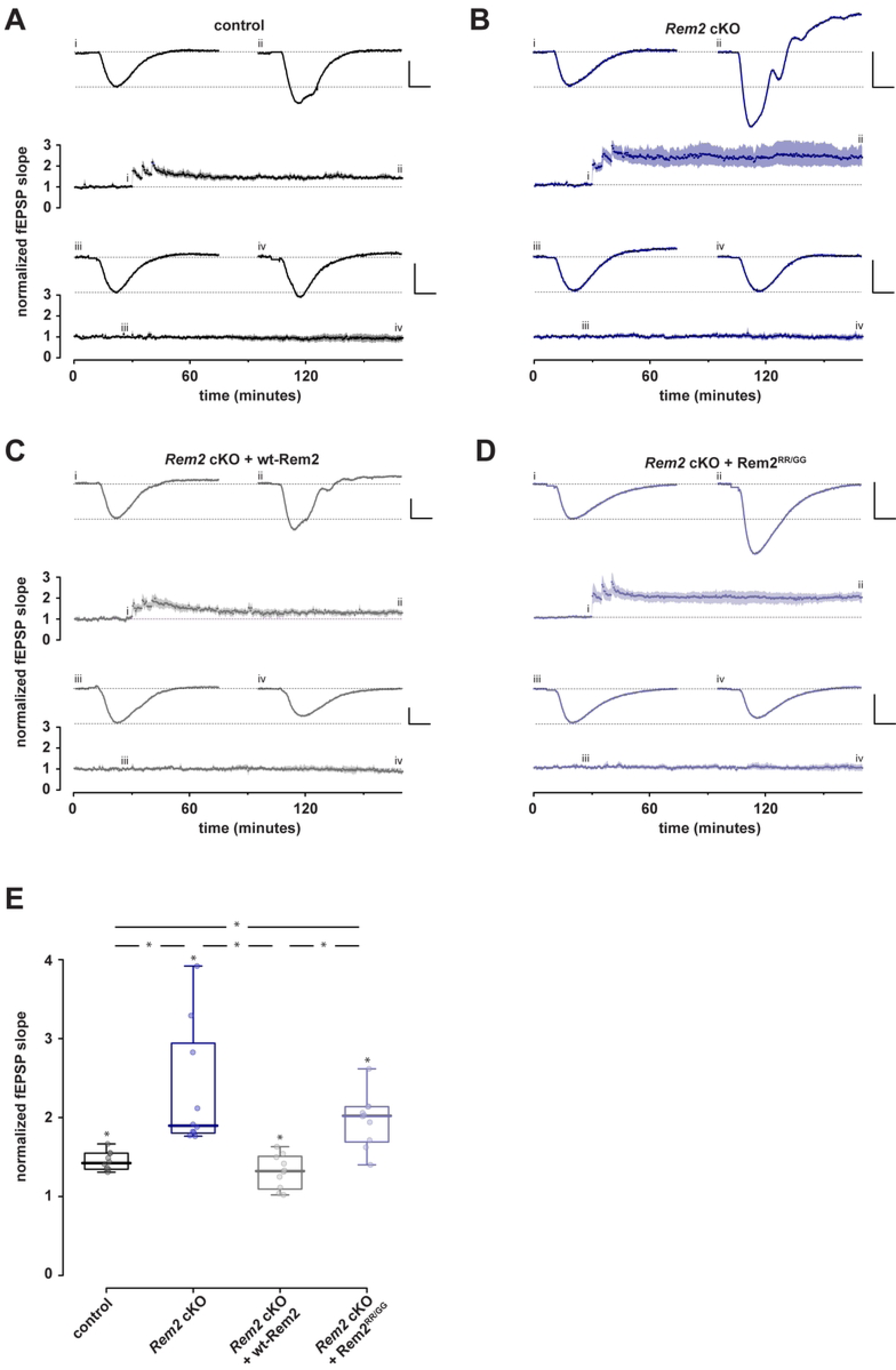
Conditional Rem2 deletion in the CA1 region of hippocampus enhances LTP at the SC-CA1 synapse. Mice from both sexes were bilaterally injected at P27-P29 with virus and acute hippocampal slices were prepared from injected mice for field recordings at P42-P46. Following a stable baseline period at least 30 minutes, LTP was induced using three high-frequency stimulation bursts at 100 Hz. Comparisons were made at *t =*170 min. A-D). Normalised fEPSP slope is plotted as a function of time for each of 4 conditions; both test and control inputs are illustrated: (A) control, (B) *Rem2* cKO, (C) *Rem2* cKO + Rem2-wt, and (D) and *Rem2* cKO + Rem2^RR/GG^; (each *n* =10). Shaded areas are the (bootstrapped) SEM. Single examples illustrate typical fEPSPs taken at the time points indicated (i, ii, iii, and iv). Vertical and horizontal scale lines indicate 0.5 mV and 5 ms, respectively. E) Summary box plots of normalized fEPSP slopes in the 4 conditions. Any significant effect is illustrated by *.

Given that Rem2 is a potent inhibitor of CaMKII kinase activity, we sought to determine if the enhanced LTP observed in the *Rem2* cKO mice is due to increased CaMKII activity as opposed to an unknown synaptic function of Rem2 that is CaMKII-independent. To address this question, we injected 4-week-old *Rem2^flx/flx^* mice with virus expressing GFP (control) or AAV9-hSyn-CreGFP (*Rem2* cKO) and isolated acute hippocampal slices at 14 d.p.i. for hippocampal field recordings. In these recordings the CaMKII inhibitory peptide myr-CN27 [52] was bath-applied following a 30-minute baseline to block LTP induction (Figure 4). Following 50 minutes of myr-CN27 exposure, we delivered three 100Hz tetani with myr-CN27 still in bath. After delivery of the tetani myr-CN27 washout began at *t =*80 min. After 60 minutes of myr-CN27 washout, we delivered another three 100Hz tetani and then recorded fEPSPs for another 60 minutes to ascertain if LTP was recovered.

**Figure 4.**
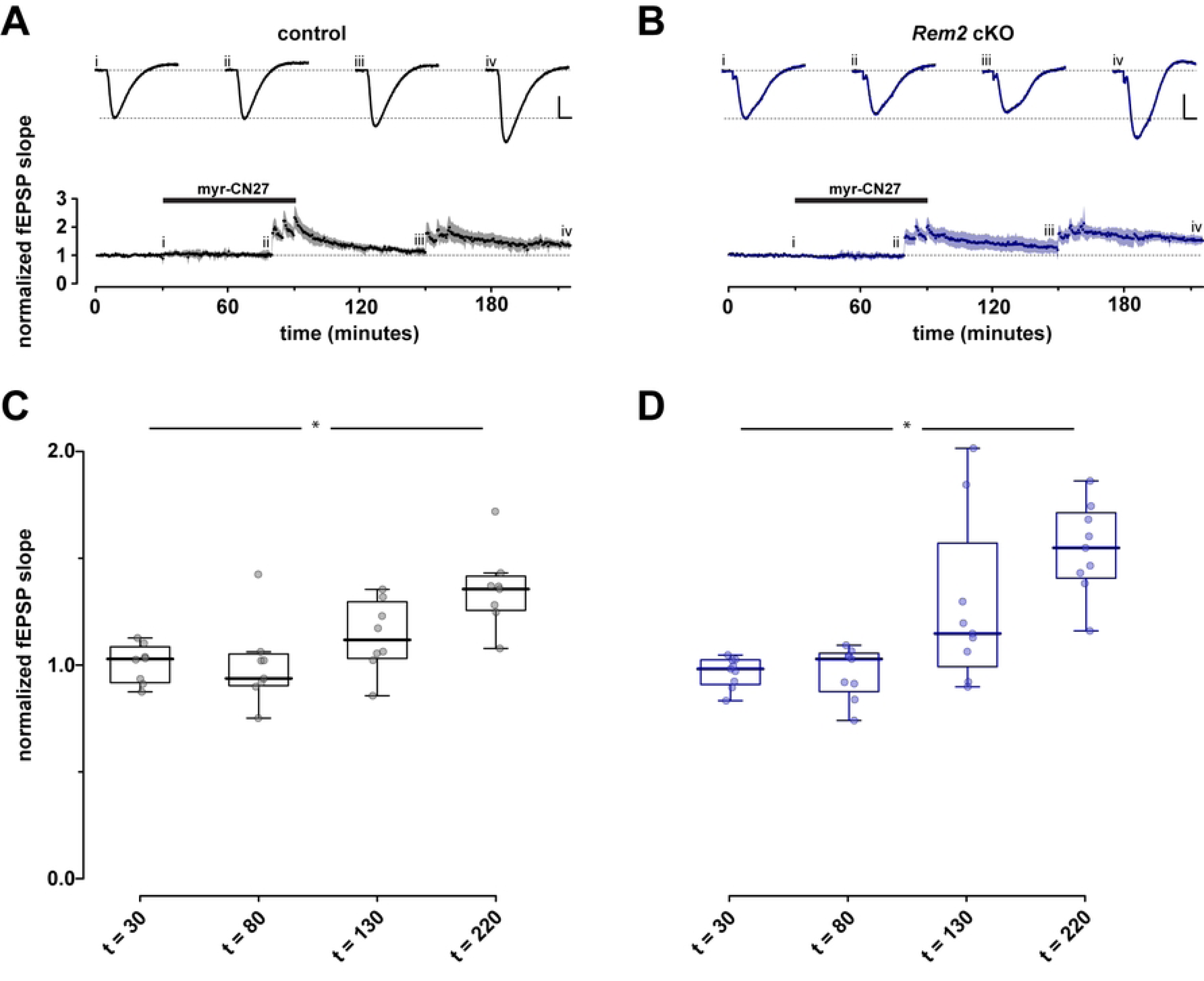
LTP observed in *Rem2* cKO is CaMKII dependent. After a stable baseline (30 min) was obtained myr-CN27 (1 µM) was bath applied for 50 minutes then three 100 Hz tetani were delivered with myr-CN27 still in bath after which myr-CN27 was washed out for 60 minutes. Subsequently three 100Hz tetani were delivered and the recording was continued for 60 minutes. A, B) Normalized fEPSP slope as a function of time for control (A, *n =* 8) and *Rem2* cKO (B, *n =* 9). Shaded areas are the (bootstrapped) SEM. Traces above the time plots represent the raw data comprising averages of time points within each experiment taken at the time points indicated (i, ii, etc.). As before, vertical and horizontal scale lines indicate 0.5 mV and 5 ms, respectively. C, D) Summary box plot to illustrate normalized fEPSP slope for control (A, *n =* 8) and *Rem2* cKO (B, *n =* 9).

Our results show that presence of myr-CN27 in bath blocks the induction of LTP in both control (Figure 4A, C, *n =* 8) and *Rem2* cKO conditions (Figure 4B, D, *n =* 9), strongly suggesting that the enhanced LTP observed in the *Rem2* cKO animals is dependent on CaMKII. The ability of this synapse to undergo LTP in both control and *Rem2* cKO slices after myr-CN27 washout also supports this idea. However, we failed to observe enhanced LTP in the *Rem2* cKO slices compared to control upon myr-CN27 washout. Despite this fact, these data are consistent with the idea that enhanced LTP observed upon Rem2 knockout is CaMKII-dependent because if Rem2 were to signal through a CaMKII-independent pathway to inhibit LTP, myr-CN27 application in the *Rem2* cKO condition should still have yielded some LTP induction and expression.

Our results so far suggest a model whereby Rem2 interacts with CaMKII at synapses to limit LTP. Although a previous study reports the co-localization of Rem2-GFP and mRuby-CaMKII in dendritic spines using diffraction limited light microscopy [42], an analysis of Rem2-CaMKII dynamics at individual synapses undergoing plasticity has not been investigated. We hypothesized that CaMKII activation will cause a change in its association with Rem2 at the synapse. To address this question, we employed two-photon excitation and fluorescence lifetime imaging microscopy (2pFLIM) which uses fluorescence resonance energy transfer (FRET)-based sensors to measure protein-protein interactions in living cells [53]. Cultured hippocampal slices isolated from P6-9 wild type mouse pups were infected at DIV3 with adeno-associated virus (AAV-DJ) encoding Clover-CaMKIIα as the donor and AAV-DJ encoding mCherry-Rem2 as the acceptor. To induce structural plasticity, two-photon MNI-glutamate uncaging was carried out at 13–14DIV at 720 nm (30 trains, 0.5 Hz, 6 ms duration/pulse, 6 mW) at the tip of the spines (Figure 5A-C) as described previously [54,55]. Two-photon excitation at 1000 nm was used to excite Clover-CaMKIIα (Figure 5A-C) at the designated spines and the FRET signal was monitored using 2pFLIM.

**Figure 5.**
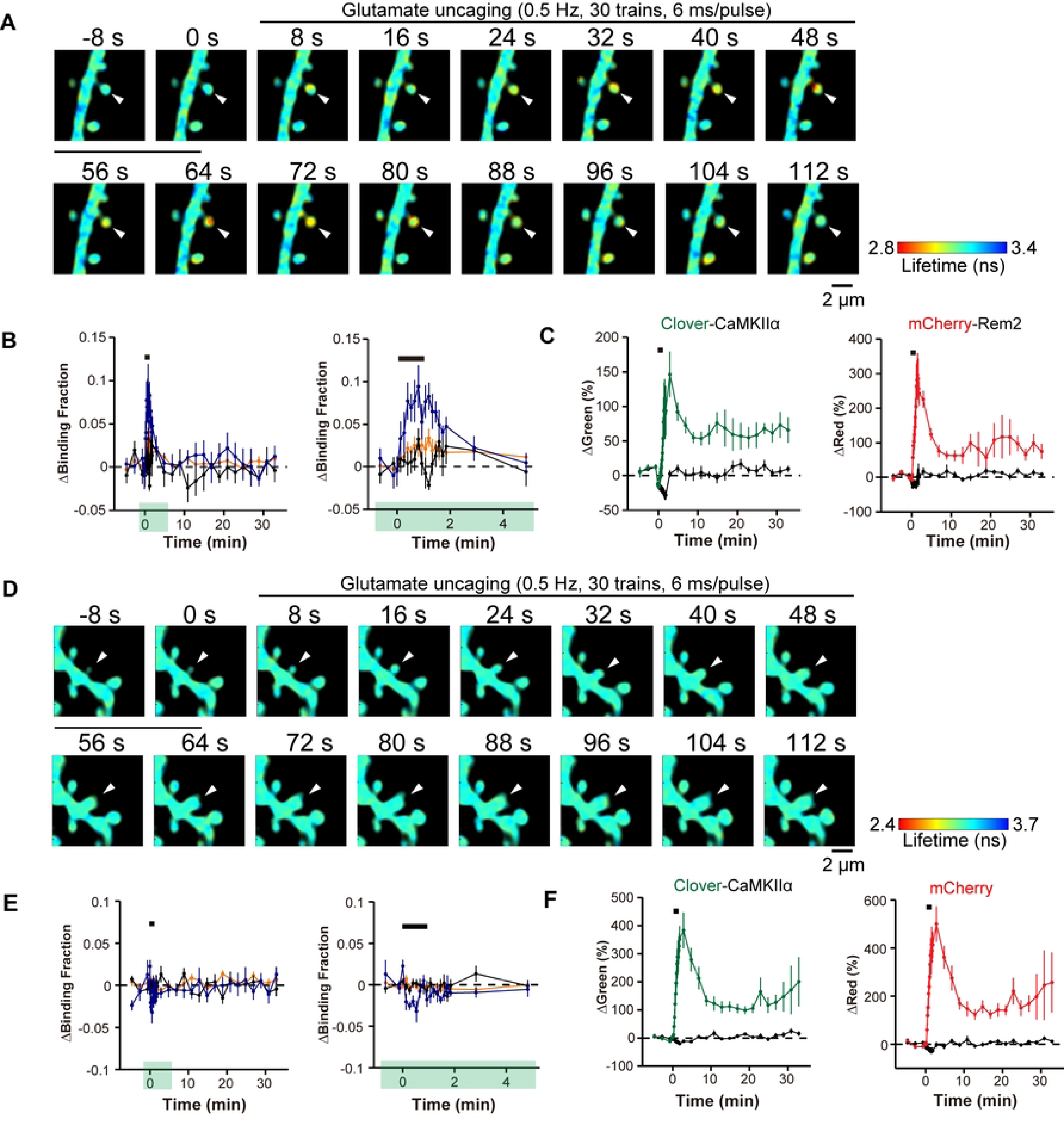
Rem2 interacts with CaMKII in a 2-photon FLIM-FRET assay. Representative fluorescence lifetime images of Clover-CaMKIIα and mCherry-Rem2 interaction (A) or Clover-CaMKIIα and mCherry interaction (D) during spine structural plasticity induced by two-photon glutamate uncaging. The arrowhead indicates the stimulated spine. Warmer colors indicate shorter fluorescent lifetimes, therefore increased interaction between proteins. B) Time course of CaMKII and Rem2 interaction measured as a change in the fraction of Clover-CaMKIIα bound to mCherry-Rem2 in stimulated spines (blue, *n =* 7), the dendritic shaft beside the stimulated spines (orange, *n =* 7), and adjacent spines (black, *n =* 7). The right panel shows a 5-minute window of the imaging data from the left panel as indicated by green highlighting. Error bars = SEM. *n =* 5 neurons. The black bar indicates glutamate uncaging. C) Averaged time course of spine volume change in the same experiments as in B for Clover-CaMKIIα and mCherry-Rem2 in stimulated spines (green for Clover-CaMKIIα, red for mCherry-Rem2, *n =* 7) and in adjacent spines (black, *n =* 7). E) Time course of CaMKII and mCherry interaction measured as a change in the fraction of Clover-CaMKIIα bound to mCherry in stimulated spines (blue, *n =*7), the dendritic shaft beside the stimulated spines (orange, *n =* 7), and adjacent spines (black, *n =* 14). The right panel shows a 5-minute window of the imaging data from the left panel as indicated by green highlighting. Error bars = SEM. *n =* 6 neurons. F) Averaged time course of spine volume change in the same experiments as in E for Clover-CaMKIIα and mCherry, in stimulated spines (green for Clover-CaMKIIα, red for mCherry, *n =* 7) and in adjacent spines (black, *n =* 7). The black bar indicates glutamate uncaging.

Upon induction of spine structural plasticity, Clover-CaMKIIα and mCherry-Rem2 demonstrated binding within 30 s in the stimulated spines which decayed over the course of two minutes (Figure 5B). We monitored the change in spine volume in response to glutamate uncaging using both the red and green signals to confirm that structural plasticity had occurred (Figure 5C). In addition, we failed to observe binding between Clover-CaMKIIα and mCherry alone, indicating that the observed change in Clover-CaMKIIα binding requires the presence of Rem2. Taken together, these data demonstrate that CaMKII and Rem2 interact in dendritic spines during structural plasticity.

## Discussion

CaMKII is a well-characterized, abundant protein kinase that mediates important functions in diverse cell types. In the nervous system, activation of CaMKII is both necessary and sufficient for the induction of NMDAR-dependent LTP, a biological correlate of learning and memory, at adult synapses in the hippocampus [23]. The signaling pathways downstream of CaMKII which lead to synaptic potentiation are well-established [1,3,23]. Automonous CaMKII is not maximally active under basal conditions, and its activity can be further stimulated ∼5-fold by subsequent Ca^2+^/CaM [56] indicating a need for an endogenous CaMKII inhibitor to constrain LTP. The role of endogenous, negative regulators of CaMKII activity, which could function to impede runaway potentiation, is not well understood. In this study we sought to determine if Rem2 functions to regulate LTP via its interaction with CaMKII. This line of inquiry was based upon a previous proteomic analysis that identified all 4 isozymes of CaMKII as interacting proteins with Rem2 and led us to the unexpected discovery that Rem2 is a direct, endogenous inhibitor of CaMKIIα activity *in vitro* [24].

Here we provide the following evidence demonstrating that Rem2 acts as a post-synaptic brake on CaMKII-dependent LTP. First, we demonstrated that knockout of *Rem2* in CA1 pyramidal neurons results in enhanced LTP at the SC-CA1 synapse. This effect was rescued with reintroduction of wild-type Rem2 but not with a *Rem2* mutant that fails to inhibit CaMKII *in vitro*. Next, we showed that the enhanced LTP observed upon *Rem2* knockout is dependent on CaMKII as the specific CaMKII inhibitor myr-CN27 blocks this effect. Lastly, we observed an activity-dependent interaction between CaMKIIα and Rem2 at glutamatergic synapses in dendritic spines in the context of structural plasticity. Interestingly, under the strong multiple stimulus condition employed in this study (Figs. 3, 4), LTP was thought to be saturated at the SC-CA1 synapse, i.e. potentiation could not be increased by further stimulation, presumably due to limitations on the components of LTP expression such as AMPAR recruitment to the synapse. Our novel and unexpected finding that Rem2 deletion enhances LTP strongly suggests that LTP was not in fact saturated in earlier studies because Rem2, a previously unappreciated negative regulator of CaMKII-dependent LTP, was acting to restrict LTP at this synapse.

LTP induction by CaMKII involves both a catalytic (i.e. phosphorylation at T286) and a structural, non-catalytic function (i.e. GluN2B binding) for CaMKII in LTP induction [20,21]. Previous work from our lab demonstrated that Rem2 does not inhibit CaMKII autophosphorylation on T286 but does inhibit the autonomously active form of CaMKII [24]. Therefore, what is the connection between Rem2 inhibition of CaMKII catalytic activity and LTP *in vivo*, given the fact that Rem2^RR/GG^ fails to rescue the enhanced LTP observed with Rem2 knockout? We posit that once CaMKII is autonomously active and the catalytic domain is persistently exposed, the N-terminus of Rem2 accesses the substrate binding site, providing block of ongoing autonomous activity to exogenous substrates and GluN2B binding resulting in inhibition of CaMKII-dependent LTP. Note that while we previously demonstrated that Rem2^RR/GG^ fails to inhibit CaMKII in vitro [24], we do not yet know if or how these amino acid substitutions affect the ability of Rem2 to bind to CaMKII. One possibility is that the RR→GG substitutions alter the affinity of the Rem2 N-terminus for the substrate binding region on CaMKII resulting in a lack of association between Rem2 and CaMKII and thereby relieving any Rem2 interference in GluN2B binding to CaMKII. Thus, although our initial logic for testing the ability of Rem2^RR/GG^ to rescue enhanced LTP was based upon a purely catalytic model of CaMKII function, it is possible that Rem2 also inhibits the structural function of CaMKII in LTP induction.

How might Rem2 interaction with CaMKII fit into our current understanding of CaMKII regulation and function at the synapse? Of course, Rem2 cannot entirely block CaMKII function or else LTP would never be achieved. It could be that Rem2 is an alternative binding partner to GluN2B for CaMKII. In this scenario, Rem2:CaMKII could sequester a subset of active CaMKII from the synapse while also inhibiting its catalytic activity, thereby blocking further LTP expression. In fact, our 2pFLIM-FRET data is consistent with Rem2 sequestration of CaMKII out of the spine head upon binding (Fig. 5). In parallel GluN2B:CaMKII could localize a distinct subset of active CaMKII to the postsynaptic density allowing for persistent LTP at the synapse. While the stoichiometry of Rem2:CaMKII and GluN2B:CaMKII has yet to be determined, CaMKII is approximately 100x more concentrated that GluN2B at the synapse [13] and given its abundance, CaMKII could be present in excess of Rem2 protein also. A difference in relative abundance between Rem2 and CaMKII at synapses could also explain why we failed to observe enhanced LTP in the *Rem2* cKO slices compared to control upon myr-CN27 washout (Fig. 4). If myr-CN27 washout was incomplete, the remaining myr-CN27 bound to CaMKII could obscure the absence of Rem2 inhibition at the synapse by continuing to provide a moderate amount of CaMKII inhibition.

Previous studies from our lab demonstrated that Rem2 is also a CaMKII substrate and is phosphorylated on 4 conserved sites by CaMKII [38]. We further speculate that Rem2 inhibition of CaMKII is overcome by the eventual phosphorylation of Rem2 by CaMKII. Phosphorylation of Rem2 by CaMKII could cause the dissociation of Rem2 from CaMKII holoenzyme, thus serving as a negative feedback loop for Rem2 inhibition of CaMKII. This mechanism would effectively provide a time limit on Rem2 inhibition as potent, lasting inhibition could have dire effects on neuronal signaling.

In addition to Rem2, only the 79 amino acid CaMKIIN isoforms α and δ [57,58] and the postsynaptic density protein Densin-180 [59] are potential endogenous inhibitors of CaMKII in mammals. Although peptides based upon CaMKIIN sequences are widely used to inhibit CaMKII and LTP in vitro and ex vivo, the function of CaMKIIN isoforms in regulating CaMKII-dependent synaptic plasticity is not well explored, although CaMKIIN1 has been shown to play a role in long-term memory after retrieval in dorsal hippocampus [60]. Interestingly, deletion of the Densin-180 gene interferes with localization of key proteins such as α-actinin to the PSD and impairs Long-term Depression (LTD) at the SC-CA1 synapse while having no effect on LTP [61]. It is unclear which if any of these effects are dependent upon Densin-180 inhibition of CaMKII; it has been suggested that the role of Densin-180 in regulating voltage-gated calcium channel trafficking could underlie its role in synaptic plasticity [62].

Our results represent a paradigm shift in the understanding of LTP. Hundreds of structural, biochemical and functional studies inform the current model of CaMKII activation that underlies SC-CA1 LTP with little consideration of endogenous CaMKII inhibitors such as Rem2. Our novel and unexpected finding that Rem2 deletion enhances LTP reveals an unsuspected negative regulation on CaMKII that limits LTP and demonstrates that the magnitude of LTP can be increased at this synapse, shedding new light on our thinking about CaMKII regulation of LTP.

## Acknowledgements

We thank Dr. Leslie Griffith, Dr. Michael Marr and the members of the Paradis and Murakoshi labs for comments and suggestions throughout the project. This work was supported by National Institutes of Health grant R01NS065856 (S.P.), JSPS KAKENHI grant numbers 22H02724/22H05549/23H04244 (H.M.), Frontier Photonic Sciences Project of National Institutes of Natural Sciences (NINS)(H.M.).

## References

1. Lisman J, Schulman H, Cline H. The molecular basis of CaMKII function in synaptic and behavioural memory. Nat Rev Neurosci. 2002;3: 175–190. doi:10.1038/nrn753

2. Sanhueza M, Fernandez-Villalobos G, Stein IS, Kasumova G, Zhang P, Bayer KU, et al. Role of the CaMKII/NMDA Receptor Complex in the Maintenance of Synaptic Strength. J Neurosci. 2011;31: 9170– 9178. doi:10.1523/JNEUROSCI.1250-11.2011

3. Lisman J, Yasuda R, Raghavachari S. Mechanisms of CaMKII action in long-term potentiation. Nat Rev Neurosci. 2012;13: 169–182. doi:10.1038/nrn3192

4. Bhattacharyya M, Karandur D, Kuriyan J. Structural insights into the regulation of Ca2+ /calmodulin-dependent protein kinase II (Camkii). Cold Spring Harb Perspect Biol. 2020;12: 1–20. doi:10.1101/cshperspect.a035147

5. Kovalchuk Y, Eilers J, Lisman J, Konnerth A. NMDA Receptor-Mediated Subthreshold Ca ^2+^ Signals in Spines of Hippocampal Neurons. J Neurosci. 2000;20: 1791–1799. doi:10.1523/JNEUROSCI.20-05-01791.2000

6. Hoffman L, Stein RA, Colbran RJ, Mchaourab HS. Conformational changes underlying calcium/calmodulin-dependent protein kinase II activation: Conformational changes underlying CaMKII activation. EMBO J. 2011;30: 1251–1262. doi:10.1038/emboj.2011.40

7. Giese KP, Fedorov NB, Filipkowski RK, Silva AJ. Autophosphorylation at Thr286 of the α calcium-calmodulin kinase II in LTP and learning. Science. 1998;279: 870–873. doi:10.1126/science.279.5352.870

8. Shi S-H, Hayashi Y, Petralia RS, Zaman SH, Wenthold RJ, Svoboda K, et al. Rapid Spine Delivery and Redistribution of AMPA Receptors After Synaptic NMDA Receptor Activation. Science. 1999;284: 1811– 1816. doi:10.1126/science.284.5421.1811

9. Derkach V, Barria A, Soderling TR. Ca ^2+^ /calmodulin-kinase II enhances channel conductance of α-amino-3-hydroxy-5-methyl-4-isoxazolepropionate type glutamate receptors. Proc Natl Acad Sci. 1999;96: 3269–3274. doi:10.1073/pnas.96.6.3269

10. Park J, Chávez AE, Mineur YS, Morimoto-Tomita M, Lutzu S, Kim KS, et al. CaMKII Phosphorylation of TARPγ-8 Is a Mediator of LTP and Learning and Memory. Neuron. 2016;92: 75–83. doi:10.1016/j.neuron.2016.09.002

11. Xie Z, Srivastava DP, Photowala H, Kai L, Cahill ME, Woolfrey KM, et al. Kalirin-7 Controls Activity-Dependent Structural and Functional Plasticity of Dendritic Spines. Neuron. 2007;56: 640–656. doi:10.1016/j.neuron.2007.10.005

12. Herring BE, Nicoll RA. Kalirin and Trio proteins serve critical roles in excitatory synaptic transmission and LTP. Proc Natl Acad Sci. 2016;113: 2264–2269. doi:10.1073/pnas.1600179113

13. Yasuda R, Hayashi Y, Hell JW. CaMKII: a central molecular organizer of synaptic plasticity, learning and memory. Nat Rev Neurosci. 2022;23: 666–682. doi:10.1038/s41583-022-00624-2

14. Vest RS, Davies KD, O’Leary H, Port JD, Bayer KU. Dual mechanism of a natural CaMKII inhibitor. Mol Biol Cell. 2007;18: 5024–5033. doi:10.1091/mbc.e07-02-0185

15. Incontro S, Díaz-Alonso J, Iafrati J, Vieira M, Asensio CS, Sohal VS, et al. The CaMKII/NMDA receptor complex controls hippocampal synaptic transmission by kinase-dependent and independent mechanisms. Nat Commun. 2018;9: 2069. doi:10.1038/s41467-018-04439-7

16. Strack S, Colbran RJ. Autophosphorylation-dependent Targeting of Calcium/ Calmodulin-dependent Protein Kinase II by the NR2B Subunit of theN-Methyl-d-aspartate Receptor. J Biol Chem. 1998;273: 20689–20692. doi:10.1074/jbc.273.33.20689

17. Leonard AS, Lim IA, Hemsworth DE, Horne MC, Hell JW. Calcium/calmodulin-dependent protein kinase II is associated with the N-methyl-D-aspartate receptor. Proc Natl Acad Sci U S A. 1999;96: 3239–3244. doi:10.1073/pnas.96.6.3239

18. Barria A, Malinow R. NMDA receptor subunit composition controls synaptic plasticity by regulating binding to CaMKII. Neuron. 2005;48: 289–301. doi:10.1016/j.neuron.2005.08.034

19. Halt AR, Dallapiazza RF, Zhou Y, Stein IS, Qian H, Juntti S, et al. CaMKII binding to GluN2B is critical during memory consolidation. EMBO J. 2012;31: 1203–1216. doi:10.1038/emboj.2011.482

20. Tullis JE, Larsen ME, Rumian NL, Freund RK, Boxer EE, Brown CN, et al. LTP induction by structural rather than enzymatic functions of CaMKII. Nature. 2023;621: 146–153. doi:10.1038/s41586-023-06465-y

21. Chen X, Cai Q, Zhou J, Pleasure SJ, Schulman H, Zhang M, et al. CaMKII autophosphorylation but not downstream kinase activity is required for synaptic memory. BioRxiv Prepr Serv Biol. 2023. doi:10.1101/2023.08.25.554912

22. Özden C, Sloutsky R, Mitsugi T, Santos N, Agnello E, Gaubitz C, et al. CaMKII binds both substrates and activators at the active site. Cell Rep. 2022;40. doi:10.1016/j.celrep.2022.111064

23. Nicoll RA, Schulman H. Synaptic memory and CaMKII. Physiol Rev. 2023;103: 2897–2945. doi:10.1152/physrev.00034.2022

24. Royer L, Herzog JJ, Kenny K, Tzvetkova B, Cochrane JC, Marr MT, et al. The Ras-like GTPase Rem2 is a potent inhibitor of calcium/calmodulin-dependent kinase II activity. J Biol Chem. 2018;293: 14798– 14811. doi:10.1074/jbc.RA118.003560

25. Finlin BS, Shao H, Kadono-Okuda K, Guo N, Andres DA. Rem2, a new member of the Rem/Rad/Gem/Kir family of Ras-related GTPases. Biochem J. 2000;347: 223. doi:10.1042/0264-6021:3470223

26. Reymond P, Coquard A, Chenon M, Zeghouf M, Marjou AE, Thompson A, et al. Structure of the GDP-bound G domain of the RGK protein Rem2. Acta Crystallograph Sect F Struct Biol Cryst Commun. 2012;68: 626–631. doi:10.1107/S1744309112013541

27. Sasson Y, Navon-Perry L, Huppert D, Hirsch JA. RGK Family G-Domain:GTP Analog Complex Structures and Nucleotide-Binding Properties. J Mol Biol. 2011;413: 372–389. doi:10.1016/j.jmb.2011.08.017

28. Bernal Astrain G, Nikolova M, Smith MJ. Functional diversity in the RAS subfamily of small GTPases. Biochem Soc Trans. 2022;50: 921–933. doi:10.1042/BST20211166

29. Zhu J, Tseng Y-H, Kantor JD, Rhodes CJ, Zetter BR, Moyers JS, et al. Interaction of the Ras-related protein associated with diabetes Rad and the putative tumor metastasis suppressor NM23 provides a novel mechanism of GTPase regulation. Proc Natl Acad Sci. 1999;96: 14911–14918. doi:10.1073/pnas.96.26.14911

30. Stiegler AL, Boggon TJ. The pseudoGTPase group of pseudoenzymes. FEBS J. 2020;287: 4232–4245. doi:10.1111/febs.15554

31. Correll RN, Pang C, Niedowicz DM, Finlin BS, Andres DA. The RGK family of GTP-binding proteins: Regulators of voltage-dependent calcium channels and cytoskeleton remodeling. Cell Signal. 2008;20: 292–300. doi:10.1016/j.cellsig.2007.10.028

32. Chen H, Puhl HL, Niu S-L, Mitchell DC, Ikeda SR. Expression of Rem2, an RGK Family Small GTPase, Reduces N-Type Calcium Current without Affecting Channel Surface Density. J Neurosci. 2005;25: 9762– 9772. doi:10.1523/JNEUROSCI.3111-05.2005

33. Finlin BS, Mosley AL, Crump SM, Correll RN, Özcan S, Satin J, et al. Regulation of L-type Ca2+ Channel Activity and Insulin Secretion by the Rem2 GTPase. J Biol Chem. 2005;280: 41864–41871. doi:10.1074/jbc.M414261200

34. Leyris J-P, Gondeau C, Charnet A, Delattre C, Rousset M, Cens T, et al. RGK GTPase-dependent Ca V 2.1 Ca 2+ channel inhibition is independent of Ca V β–subunit-induced current potentiation. FASEB J. 2009;23: 2627–2638. doi:10.1096/fj.08-122135

35. Moore AR, Ghiretti AE, Paradis S. A Loss-Of-Function Analysis Reveals That Endogenous Rem2 Promotes Functional Glutamatergic Synapse Formation and Restricts Dendritic Complexity. PLoS ONE. 2013;8. doi:10.1371/journal.pone.0074751

36. Paradis S, Harrar DB, Lin Y, Koon AC, Hauser JL, Griffith EC, et al. An RNAi-Based Approach Identifies Molecules Required for Glutamatergic and GABAergic Synapse Development. Neuron. 2007;53: 217–232. doi:10.1016/j.neuron.2006.12.012

37. Ghiretti AE, Paradis S. The GTPase Rem2 regulates synapse development and dendritic morphology. Dev Neurobiol. 2011;71: 374–389. doi:10.1002/dneu.20868

38. Ghiretti AE, Kenny K, Marr MT, Paradis S. CaMKII-Dependent Phosphorylation of the GTPase Rem2 Is Required to Restrict Dendritic Complexity. J Neurosci. 2013;33: 6504–6515. doi:10.1523/JNEUROSCI.3861-12.2013

39. Ghiretti AE, Moore AR, Brenner RG, Chen L-F, West AE, Lau NC, et al. Rem2 Is an Activity-Dependent Negative Regulator of Dendritic Complexity In Vivo. J Neurosci. 2014;34: 392–407. doi:10.1523/JNEUROSCI.1328-13.2014

40. Moore AR, Richards SE, Kenny K, Royer L, Chan U, Flavahan K, et al. Rem2 stabilizes intrinsic excitability and spontaneous firing in visual circuits. eLife. 2018;7. doi:10.7554/eLife.33092

41. Richards SEV, Moore AR, Nam AY, Saxena S, Paradis S, Van Hooser SD. Experience-Dependent Development of Dendritic Arbors in Mouse Visual Cortex. J Neurosci. 2020;40: 6536–6556. doi:10.1523/JNEUROSCI.2910-19.2020

42. Flynn R, Labrie-Dion E, Bernier N, Colicos MA, Koninck PD, Zamponi GW. Activity-Dependent Subcellular Cotrafficking of the Small GTPase Rem2 and Ca2+/CaM-Dependent Protein Kinase IIα. PLoS ONE. 2012;7: e41185. doi:10.1371/journal.pone.0041185

43. Anderson WW, Collingridge GL. Capabilities of the WinLTP data acquisition program extending beyond basic LTP experimental functions. J Neurosci Methods. 2007;162: 346–356. doi:10.1016/j.jneumeth.2006.12.018

44. Stoppini L, Buchs P-A, Muller D. A simple method for organotypic cultures of nervous tissue. J Neurosci Methods. 1991;37: 173–182. doi:10.1016/0165-0270(91)90128-M

45. Lock M, Alvira M, Vandenberghe LH, Samanta A, Toelen J, Debyser Z, et al. Rapid, Simple, and Versatile Manufacturing of Recombinant Adeno-Associated Viral Vectors at Scale. Hum Gene Ther. 2010;21: 1259–1271. doi:10.1089/hum.2010.055

46. Shibata ACE, Ueda HH, Eto K, Onda M, Sato A, Ohba T, et al. Photoactivatable CaMKII induces synaptic plasticity in single synapses. Nat Commun. 2021;12: 751. doi:10.1038/s41467-021-21025-6

47. Yasuda M, Mayford MR. CaMKII Activation in the Entorhinal Cortex Disrupts Previously Encoded Spatial Memory. Neuron. 2006;50: 309–318. doi:10.1016/j.neuron.2006.03.035

48. Murakoshi H. Optogenetic Imaging of Protein Activity Using Two-Photon Fluorescence Lifetime Imaging Microscopy. Advances in Experimental Medicine and Biology. 2021. pp. 295–308. doi:10.1007/978-981-15-8763-4_18

49. Pologruto TA, Sabatini BL, Svoboda K. ScanImage: Flexible software for operating laser scanning microscopes. Biomed Eng OnLine. 2003;2: 13. doi:10.1186/1475-925X-2-13

50. Lein ES, Hawrylycz MJ, Ao N, Ayres M, Bensinger A, Bernard A, et al. Genome-wide atlas of gene expression in the adult mouse brain. Nature. 2007;445: 168–176. doi:10.1038/nature05453

51. Sciences AI for B. Allen Mouse Brain Atlas [dataset]. 2004. Available: https://mouse.brain-map.org/gene/show/80094

52. Tao W, Lee J, Chen X, Diaz-Alonso J, Zhou J, Pleasure S, et al. Synaptic memory requires CaMKII. eLife. 2021;10. doi:10.7554/eLife.60360

53. Ueda HH, Nagasawa Y, Murakoshi H. Imaging intracellular protein interactions/activity in neurons using 2-photon fluorescence lifetime imaging microscopy. Neurosci Res. 2022;179: 31–38. doi:10.1016/j.neures.2021.10.004

54. Matsuzaki M, Honkura N, Ellis-Davies GCR, Kasai H. Structural basis of long-term potentiation in single dendritic spines. Nature. 2004;429: 761–766. doi:10.1038/nature02617

55. Chang J-Y, Nakahata Y, Hayano Y, Yasuda R. Mechanisms of Ca2+/calmodulin-dependent kinase II activation in single dendritic spines. Nat Commun. 2019;10: 2784. doi:10.1038/s41467-019-10694-z

56. Barcomb K, Buard I, Coultrap SJ, Kulbe JR, O’Leary H, Benke TA, et al. Autonomous CaMKII requires further stimulation by Ca ^2+^ /calmodulin for enhancing synaptic strength. FASEB J. 2014;28: 3810–3819. doi:10.1096/fj.14-250407

57. Chang BH, Mukherji S, Soderling TR. Characterization of a calmodulin kinase II inhibitor protein in brain. Proc Natl Acad Sci. 1998;95: 10890–10895. doi:10.1073/pnas.95.18.10890

58. Chang BH, Mukherji S, Soderling TR. Calcium/calmodulin-dependent protein kinase II inhibitor protein: localization of isoforms in rat brain. Neuroscience. 2001;102: 767–777. doi:10.1016/S0306-4522(00)00520-0

59. Jiao Y, Jalan-Sakrikar N, Robison AJ, Baucum AJ, Bass MA, Colbran RJ. Characterization of a Central Ca2+/Calmodulin-dependent Protein Kinase IIα/β Binding Domain in Densin That Selectively Modulates Glutamate Receptor Subunit Phosphorylation. J Biol Chem. 2011;286: 24806–24818. doi:10.1074/jbc.M110.216010

60. Vigil FA, Mizuno K, Lucchesi W, Valls-Comamala V, Giese KP. Prevention of long-term memory loss after retrieval by an endogenous CaMKII inhibitor. Sci Rep. 2017;7: 4040. doi:10.1038/s41598-017-04355-8

61. Carlisle HJ, Luong TN, Medina-Marino A, Schenker L, Khorosheva E, Indersmitten T, et al. Deletion of Densin-180 Results in Abnormal Behaviors Associated with Mental Illness and Reduces mGluR5 and DISC1 in the Postsynaptic Density Fraction. J Neurosci. 2011;31: 16194–16207. doi:10.1523/JNEUROSCI.5877-10.2011

62. Wang S, Stanika RI, Wang X, Hagen J, Kennedy MB, Obermair GJ, et al. Densin-180 Controls the Trafficking and Signaling of L-Type Voltage-Gated Cav1.2 Ca2+ Channels at Excitatory Synapses. J Neurosci Off J Soc Neurosci. 2017;37: 4679–4691. doi:10.1523/JNEUROSCI.2583-16.2017

